# A Phantom for Fluorescence Uniformity and Distortion Assessment of Near-Infrared Fluorescence Guided Surgery Systems

**DOI:** 10.1101/2025.05.08.652975

**Authors:** Emmanuel A. Mannoh, Edwin A. Robledo, Samuel S. Streeter, Ethan P. M. LaRochelle, Alberto J. Ruiz

**Affiliations:** QUEL Imaging, White River Junction, VT 05001 USA; Department of Orthopaedics, Geisel School of Medicine, Dartmouth College, Hanover, NH 03755 USA

**Keywords:** fluorescence-guided surgery, fluorescence uniformity, geometric distortion, tissue-simulating phantom, flat-field correction, fluorescence imaging

## Abstract

**Significance:** The expanding use of fluorescence in surgery necessitates standardized characterization methods to facilitate reproducibility and regulatory review of imaging devices. Current guidelines suggest the use of optical phantoms as tools to quantify optical system performance, yet measurements of uniformity and spatial accuracy or distortion remain challenging and are performed in an ad hoc manner or not collected at all.

**Aim:** This work introduces a photostable solid phantom, the Reference Uniformity and Distortion (RUD) phantom, and accompanying analysis code for characterizing fluorescence uniformity and geometric distortion. Additionally, the concept of fluorescence flat-field correction is explored using this phantom.

**Approach:** The RUD phantom was imaged on a custom fluorescence imaging device, as well as five commercial systems. The analysis code characterized uniformity and distortion in these systems. Flat-field correction was explored on the custom device by imaging solid fluorescent reference phantoms at different locations within the field of view.

**Results:** Successful characterization of the imaging systems’ uniformity and geometric distortion was achieved. Flat-fielding experiments showed that while it qualitatively improves the appearance of images, it could negatively impact quantitative analyses.

**Conclusions:** The RUD addresses the need for standardized characterization of fluorescence uniformity and geometric distortion. While fluorescence flat-field correction qualitatively enhances image uniformity, caution is advised as it may adversely affect quantitative accuracy.

## 1 Introduction

The use of fluorescence to guide surgery has expanded in recent years since the FDA’s 510(k) clearance of the SPY Imaging System in 2005 for assessing vascular perfusion using indocyanine green (ICG).^1^ In addition to the use of ICG for perfusion assessment, the FDA has also approved fluorescent agents to aid in cancer delineation and resection, such as 5-ALA for brain cancers,^2^ pafolacianine for ovarian^3^ and lung cancer,^4^ and pegulicianine for breast cancer.^5^ With these approvals has also come the proliferation of fluorescence guided surgery (FGS) imaging systems, making it important to have characterization and standardization methods to facilitate FDA review of new devices and allow cross-device comparisons.^6^

A Task Group formed by the American Association of Physicists in Medicine assembled a list of key metrics needed to characterize the performance of fluorescence imaging systems.^7^ This task group and recent draft guidance by the FDA^8^ highlight the importance of using phantoms for system characterization. Key metrics include: the sensitivity of the system to the fluorophore of interest, the sensitivity of the system to fluorescence at depth, spatial resolution, depth of field, overall fluorescence signal detection uniformity, and geometric distortion. Efforts have been made to develop solid phantoms that enable characterization of these key metrics.^9–13^ These efforts have largely focused on characterizing sensitivity to fluorophore concentration, sensitivity to fluorescence at depth, and spatial resolution. While some have addressed signal uniformity, these attempts have generally involved sparse sampling of the fluorescence response across the field of view,^12,13^ or homogeneous liquid phantoms^14^ that suffer from lack of reproducibility and shelf stability. Furthermore, to our knowledge, there has been no focus on developing phantoms or tools for characterizing distortion in FGS systems.

Fluorescence signal detection uniformity refers to how consistent a fluorescent signal appears across the field of view of the imaging system. It couples both the spatial uniformity of the excitation light source, and the detection uniformity of the camera system (referred to as relative illumination).^15^ Fluorescence uniformity is crucial to characterize when making clinical decisions based on a fluorescence image. For example, in a surgical procedure involving ICG perfusion, a blood vessel may appear well-perfused in the center of the image but poorly perfused when on the periphery of the field of view for systems with significant fluorescence non-uniformity. In such a scenario, the observed difference in fluorescence intensity does not reflect true biological function but rather stems from non-uniformity in the fluorescence signal detection of the imaging system. While FGS systems have generally been used clinically solely to enhance surgical contrast, recent introductions of semi-quantitative assessments, either assigning signal to background ratios in static images, or quantifying fluorescence intensity of dynamic flow, also highlight the importance of better fluorescence uniformity characterization.^16^

In an attempt to compensate for this non-uniformity in FGS systems, flat-field correction has been employed.^12,14^ The term “flat-field correction” comes from digital image processing and involves accounting for spatial non-uniformity in brightness caused by variations in sensor pixel output, imaging optics, and the effects of illumination inhomogeneities.^17,18^ While this is an established method in conventional/white light imaging, fluorescence imaging conflates excitation intensity, fluorophore emission, background signals (e.g. autofluorescence) and detector noise into a single measurement, unlike the separable illumination/reflectance in white light imaging.

Geometric distortion refers to a spatially-dependent change in magnification in the image that can alter the perceived shape and size of tissue structures.^7^ Characterizing distortion in FGS systems is equally important, as it can cause inaccuracies when estimating the size and shape of anatomical structures or their positions relative to each other.

In this manuscript, we present a single photostable solid fluorescence phantom, designed to provide assessment of near-infrared (NIR) fluorescence uniformity and geometric distortion. The use of the phantom is demonstrated in custom and commercial fluorescence imaging systems, using open-source analysis code.^19^ Finally, we explore the use of flat-field correction for fluorescence images.

## 2 Methods

### 2.1 Description of the phantom

The reference uniformity and distortion (RUD) phantom/target (SKU: RUD_STD_S800-01_TSR-01_r0, QUEL Imaging) is shown in **Figure 1**. It is designed for the simultaneous assessment of fluorescence uniformity and geometric distortion. The RUD phantom consists of a black 3D-printed light-absorbing mold back-filled with a photostable luminescent material. The luminescent material (S800-01, QUEL Imaging) is broadly excitable in the wavelength range of about 400-790 nm, with a broad emission peaking around 820 nm. The fluorescence excitation and emission spectra of the luminescent material are shown in **Supplementary Figure S1**. The top surface of the mold contains a grid of evenly spaced wells that enables the manufacturing of fluorescent dots flush with the top surface of the phantom. The fluorescent wells are each 1 mm in diameter and are spaced at center-to-center intervals of 2 mm over a 100×100 mm imaging area (2601 wells total). The overall dimensions of the target are 110 mm x 110 mm x 20 mm.

**Figure 1.**
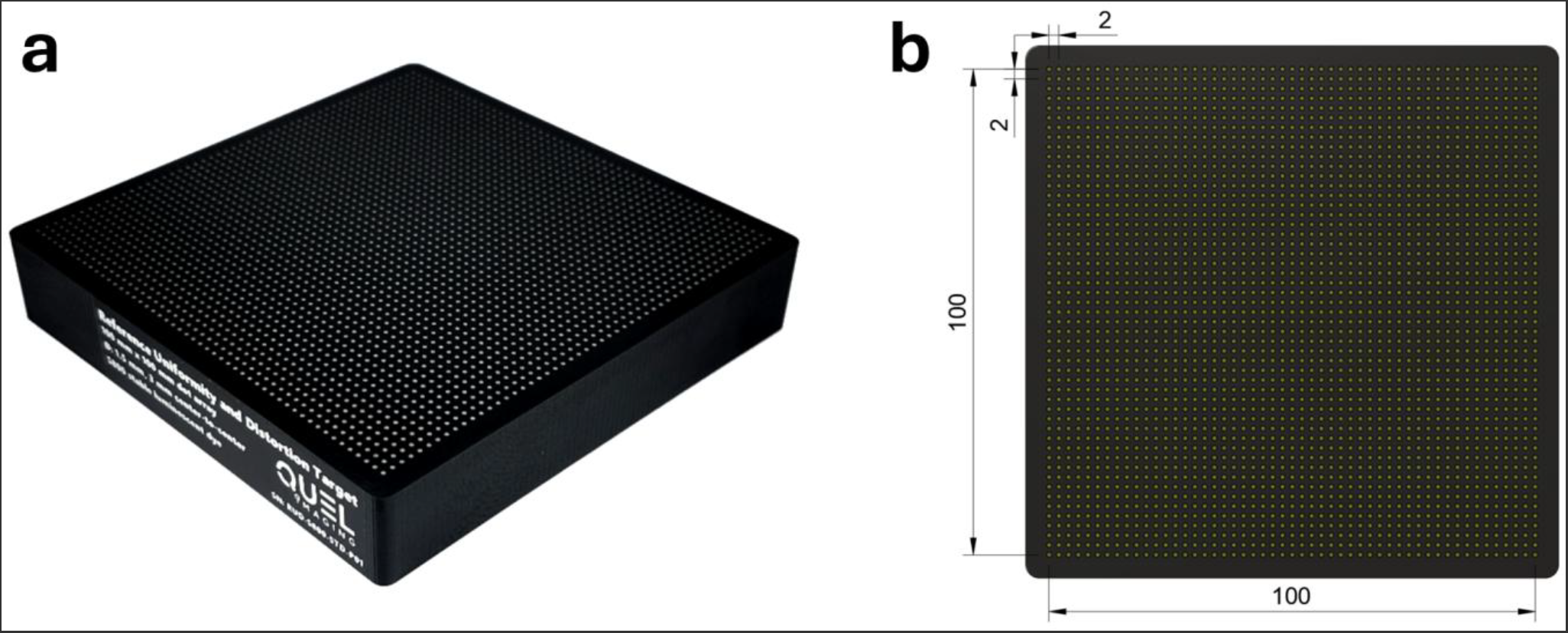
(a) Isometric white-light image of the RUD target and (b) top-down mechanical drawing showing the 100×100 mm imaging area with 1 mm diameter wells with 2 mm center-to-center spacing.

### 2.2 Obtaining fluorescence uniformity profiles with the RUD target

Custom Python code was developed to analyze RUD images, generate fluorescence uniformity maps and to quantify imaging uniformity. This code is part of an open-source library, QUEL-QAL, developed to facilitate the analysis of fluorescence reference targets,^19^ available at: https://github.com/QUEL-Imaging/quel-qal (PyPI: quel-qal). If the imaging system field of view is larger than the 100×100 mm imaging area of the phantom, multiple images of the RUD target can be acquired to obtain a fluorescence uniformity profile that spans the field of view (see **Supplementary Figure S2** for an example where four images were captured to span a ∼110x150 mm field of view). The developed code can stitch the data from these various images together to create a uniformity map that spans the full system field of view. Additionally, precise displacements of the target within the field of view can also create subsampling of the spatial resolution of the RUD to artificially increase the virtual dot-matrix generated from the combination of multiple acquisition images.

The analysis method is briefly described here. First, the pixel locations and average intensities of the fluorescent wells are extracted from the input image(s) by thresholding and identifying regions of interest (ROIs) based on connected components. The data is then fit to a surface representation using one of two methods: bivariate b-spline interpolation, or radial basis function (RBF) interpolation. The b-spline fitting method is the default and produces a smoothed representation of fluorescence uniformity from the input data. The RBF interpolation method is slower and more susceptible to noise, but preserves higher-frequency features that the b-spline method would smooth out. The surface representation is normalized so that the maximum fluorescence intensity has a value of 1. Finally, this surface representation can be visualized in multiple ways: a 3D surface plot, a 2D intensity map, line profiles, and a 2D iso-map showing regions of the field of view that are within specified percentages of the maximum fitted fluorescence intensity. Full details on the analysis methods can be found in the QUEL-QAL wiki on GitHub.^20^ Version 0.2.5 of the code was used in analyzing data and producing figures in this study.

#### 2.2.1 Assessment of uniformity quantification

A custom fluorescence imaging system was used in imaging the RUD target, and in subsequent flat-fielding experiments. This system, from here on referred to as the QUEL Imaging Box, has multiple laser excitation wavelengths and emission collection bands. For imaging the RUD target, the excitation wavelength was 760 nm, and the emission was collected with an 805 nm long-pass filter. An example of the visualizations described in **Section 2.2** produced on this imaging system is shown in **Figure 2**.

**Figure 2.**
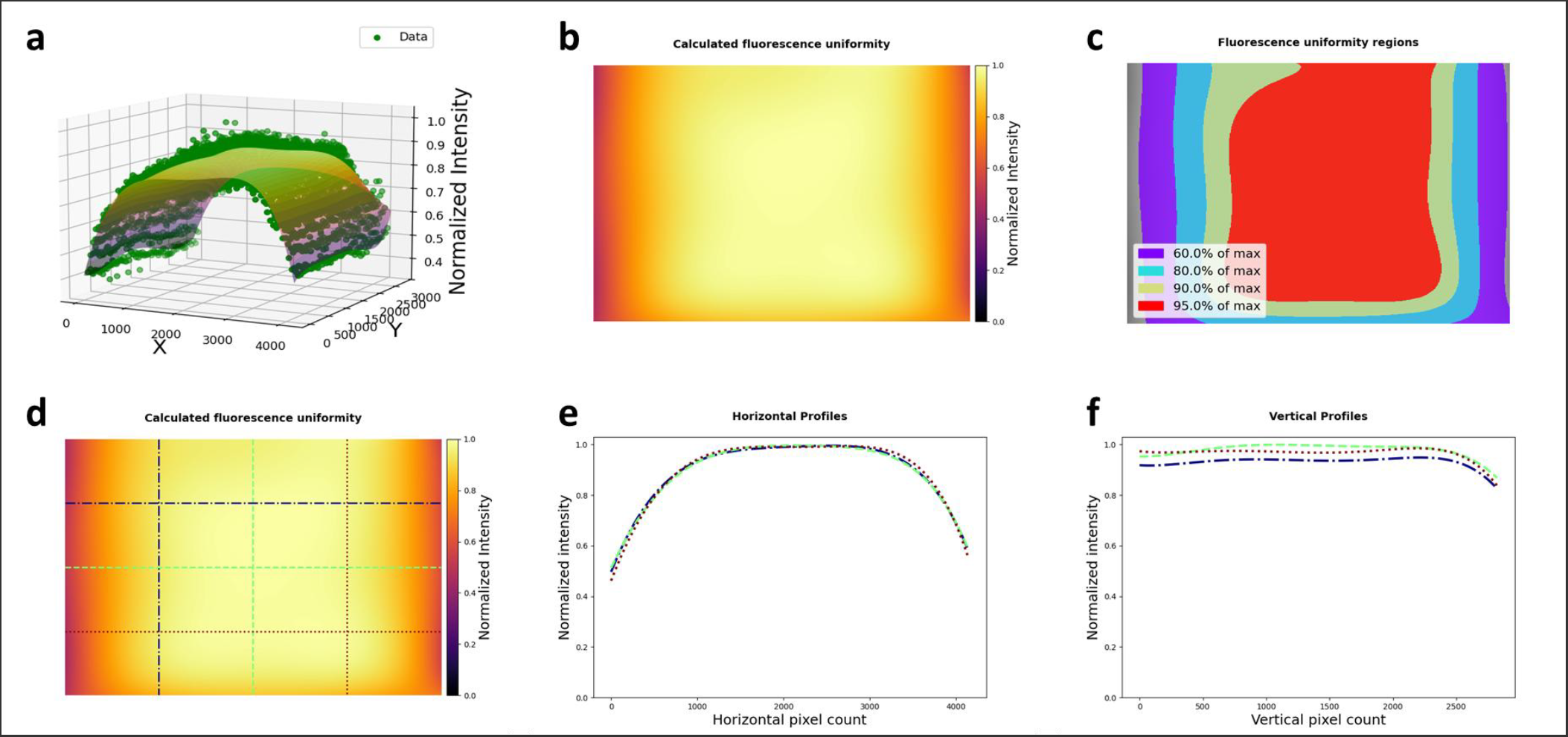
RUD analysis results using b-spline fitting method: (a) 3D plot showing extracted data and fitted surface; (b) fitted fluorescence uniformity map normalized to its maximum; (c) iso-maps, showing regions of the field of view that are at least 60%, 80%, 90%, and 95% of the maximum intensity; (d – f) line profiles across the fluorescence uniformity fit.

### 2.3 Obtaining geometric distortion with the RUD target

Custom Python code was developed to analyze images of the RUD target to quantify geometric distortion across the imaging system’s field of view. This code is part of the same open-source QUEL-QAL library used for fluorescence uniformity analysis (**Section 2.2**). Full details on the analysis can be found in the QUEL-QAL wiki.^20^ If the imaging system field-of-view exceeds the 100×100 mm imaging area of the phantom, multiple images of the RUD target can be acquired and combined to assess geometric distortion comprehensively across the full system field of view – the only caveat here is that, unlike uniformity assessment, each image must have a minimum number of fluorescent wells identified around the center of the image.

The distortion analysis method is aligned with ISO 17850:2015 – Geometric distortion (GD) measurements. Our analysis method is briefly described here. First, the pixel locations of the fluorescent wells are extracted from the image(s) using thresholding and identifying ROIs based on connected components. Next, a regular reference grid is created based on the spacing and orientation of the wells located near the center of the image. Geometric distortion is quantified by comparing the reference grid to the actual measured centroid coordinates of the fluorescent wells (**Supplementary Figure S3**). Local geometric distortion is calculated using:

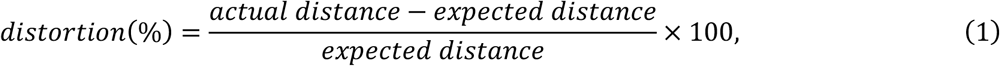

where *actual distance* is the distance from the center of the image to each fluorescent well, and *expected distance* is the distance from the center of the image to the corresponding reference grid point from the previous step. The distortion data is then plotted as a function of actual distance from the center of the field of view. In addition to this plot, the distortion data can also be used to produce a 2D map. This map effectively identifies spatial distortion patterns, such as keystone distortion resulting from object or image plane misalignment relative to the optical axis.

#### 2.3.1 Assessment of geometric distortion quantification

The QUEL Imaging Box was used to image the RUD target with the same excitation and emission parameters described in **Section 2.2.1**. An example of the distortion visualization described in **Section 2.3** produced on this system is shown in **Figure 3**. To demonstrate keystone distortion, a 3° wedge was placed beneath the RUD target and it was imaged again on the QUEL Imaging Box.

**Figure 3.**
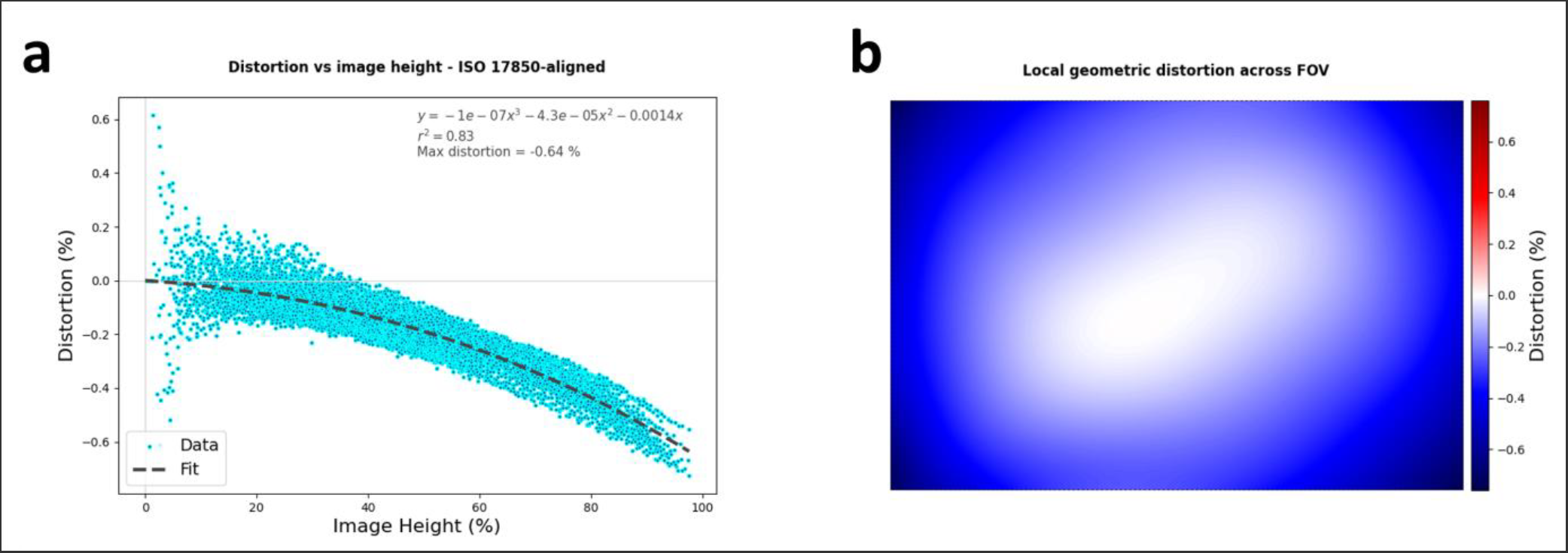
Distortion analysis results: (a) local geometric distortion as a function of image height, showing a small amount of negative or barrel distortion; (b) spatial map of distortion across the field of view, showing radial symmetry.

### 2.4 Utilizing the RUD with commercial fluorescence imaging systems

To explore the variability in fluorescence uniformity and distortion among some commercial fluorescence imaging systems, a RUD target was imaged on five commercial imaging systems at the Dartmouth Hitchcock Medical Center in Lebanon, NH. The devices were: Pentero 800 S (Zeiss, Oberkochen, Germany), Kinevo 900 S (Zeiss, Oberkochen, Germany), Modus X (Synaptive Medical, Toronto, Ontario, Canada), Pearl Impulse (LI-COR Biosciences, Lincoln, NE), and SPY-PHI (Stryker, Portage, MI). All systems are state-of-the-art, clinical fluorescence guided surgery systems except for the Pearl Impulse, which is a preclinical system. Pearl Impulse imaging data was collected using both the 700 nm and 800 nm fluorescence channels. The imaging systems are hereafter referred to as System 1, System 2, System 3, System 4A (700-channel), System 4B (800-channel), and System 5, respectively. All systems were capable of imaging the RUD target except for System 5, which has an excitation wavelength of 805 nm,^21^ which is beyond the excitation range for the RUD (**Supplementary Figure S1**). Therefore, it was excluded from further analysis. In all cases, imaging was performed with the room lights off to prevent inadvertent excitation of the RUD target by ambient light. (This measure was not required for System 4 since it is a closed-field imager.) For systems that had simultaneous white light and fluorescence imaging, white light illumination was disabled so that the results obtained would truly be representative of just the fluorescence uniformity. This was not possible for Systems 1 and 2, and so imaging was performed with the white light illumination. Each imager was aligned so that its optical axis was orthogonal to the top surface of the RUD target. If a system had zoom capabilities, imaging was performed at the lowest zoom setting to maximize field of view. For each system, as many images as needed were acquired to span the entire field of view. This required two images each for Systems 1 and 2, and one each for the rest. QUEL-QAL code (version 0.2.5) was used to quantify image uniformity and distortion for each system.

### 2.5 Flat-field correction: RUD target

To explore the possibility of using the RUD target for fluorescence flat-field correction, images of the target were first acquired with the QUEL Imaging Box and used to generate a fluorescence uniformity profile. The target was imaged in four locations across the field of view, and at each location, the target was rotated three times by 90° (for a total of four imaged orientations). So, in total, sixteen images were acquired. The images were analyzed using the analysis method described in **Section 2.2**. For the surface fitting portion, the RBF interpolation method was used, since the QUEL Imaging Box has a very structured illumination pattern on the edges of the field of view. After obtaining the fluorescence uniformity profile, the input images were each scaled by the inverse of this profile to produce “flat-field corrected” images. Then, to evaluate the performance of the correction, these flat-field corrected images were passed through the analysis algorithm again, this time using the b-spline method to avoid any high frequency features or noise. Ideally, the resulting surface should be perfectly flat.

A second set of four images was acquired with the RUD target in only one orientation and spanning the field of view of the QUEL Imaging Box. The purpose of this step was to evaluate flat-field correction on a new set of data that was not used in generating the uniformity profile for the QUEL Imaging Box. These images were scaled by the inverse of the uniformity profile generated above, and then passed through the analysis algorithm again using the b-spline method. Ideally, the resulting profile should also be flat.

### 2.6 Flat-field correction: fluorescent cylinders

For an initial investigation of fluorescence flat-field correction, three 3D-printed cylinders (Ø 10 mm) with different concentrations of ICG equivalent fluorophore – 100 nM, 3 nM, 0 nM – were imaged using the QUEL Imaging Box. The tissue-mimicking cylinders had absorption and reduced scattering of 0.021 mm-^1^ and 0.27 mm^-1^, respectively, at 800 nm. The manufacturing techniques for these 3D-printed cylinders are described in Ruiz et. al^22^. A custom holder was used to hold the cylinders next to each other and prevent fluorescence bleed-through between them. They were imaged in the following manner. For the first image, the 100 nM cylinder was placed next to the 0 nM cylinder in the center of the field of view. For the second image, these two cylinders were imaged on the left edge of the field of view, where the illumination intensity was known to drop. For the third image, the 3 nM cylinder was placed in the holder next to the 0 nM cylinder in the center of the field of view. For the fourth and final image, the 0 nM and 3 nM cylinders were imaged on the left edge of the field of view where the illumination intensity drops.

The average intensity of each cylinder in each image was obtained from a circular ROI that was about two-thirds the radius of each cylinder. All four images were then scaled by the inverse of the fluorescence uniformity profile generated from **Section 2.5**, and the average intensity of each cylinder in each image was measured again. Ideally, after flat-field correction, the intensity of each well should be consistent, regardless of its location in the field of view (i.e., on the edge of the field of view or in the center of the field of view).

### 2.7 Flat-field correction: concentration sensitivity target

A reference concentration sensitivity (RCS) target (SKU: RCS-ICG-ST01-QUEL03, QUEL Imaging) was used in this experiment. The target consists of nine wells with varying concentrations of fluorophore (in this case, ICG-equivalent dye), and serves as a dilution series to probe the fluorescence sensitivity of an imaging system.^9^ Here, a total of five images were acquired with this target. First, it was imaged in the center of the field of view, oriented such that the highest concentration well was in the top left. Then it was imaged on the left edge of the field of view where the illumination intensity drops, in four orientations rotated 90° from each other.

The images were analyzed using QUEL-QAL, with the key output metric being the linearity of the intensity-versus-fluorophore-concentration curve. Full details on the analysis methods can be found in the QUEL-QAL wiki.^23^ Briefly, the analysis method quantifies the average intensity of each well using an ROI that is half the diameter of the well, baselines by subtracting the intensity of the control (0 nM) well, normalizes to the maximum, and then fits the resulting data to the equation:

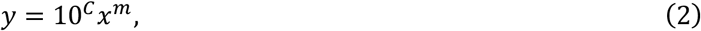

where *y* is the measured ROI fluorescence intensity, *C* is a constant, *x* is the fluorophore concentration, and *m* is a measure of linearity whose value should ideally be equal to 1.^24^

The images were then scaled by the inverse of the fluorescence uniformity profile from **Section 2.5** and reanalyzed. If the fluorescence flat-field correction is accurate, the value of *m* in the corrected images should be closer to 1 than in the uncorrected images.

## 3 Results

### 3.1 Fluorescence uniformity profiles

Four images acquired on the QUEL Imaging Box were used as input to the custom Python code for assessing fluorescence uniformity. The analysis results for this system, using b-spline fitting, are shown in **Figure 2. Figure 2a** shows a 3D plot of the normalized intensity data extracted from the images of the RUD target (green dots), and the fitted surface profile. **Figure 2b** is the fitted surface profile shown as an intensity map and normalized to its maximum. **Figure 2c** displays iso-maps, showing regions of the field of view that have intensity of at least 60%, 80%, 90%, and 95% of the maximum intensity in the fitted uniformity profile. **Figure 2d-f** show horizontal and vertical line profiles taken at evenly spaced intervals (in this case, quarters) across the field of view. **Supplementary Figure S4** shows the same data analyzed using the RBF interpolation method. This RBF analysis result highlights a notable transient dip in intensity at the outer edges of the illumination pattern that was not identified with the b-spline method.

### 3.2 Local geometric distortion

Geometric distortion was quantified using the same images of the RUD target acquired with the QUEL Imaging Box (**Section 3.1**) as input to the custom Python code. The results are shown in **Figure 3**. The calculated local geometric distortion becomes more negative with increasing image height (i.e., distance from the center of the field of view), indicating barrel distortion in this system (**Figure 3a**). The magnitude of the maximum distortion, however, is less than 1%. In **Figure 3b**, the calculated distortion is displayed as a function of the location of each data point within the field of view. The rotational symmetry of this figure suggests there is minimal, if any, keystone distortion present in this imaging system/setup.

In contrast, when a 3° wedge was placed beneath the RUD target prior to being imaged on the same system, the spatial distortion map showed an asymmetry such that the left side of the image had positive distortion, while the right side of the image had negative distortion (**Supplementary Figure S5b**). Additionally, the distortion-versus-image-height plot (**Supplementary Figure S5a**) shows two groups of data points, one with slightly positive distortion, and the other with more negative distortion.

### 3.3 Fluorescence uniformity and distortion of commercial devices

The results of fluorescence uniformity and distortion assessments for the commercial imaging systems are shown in **Figure 4** and **Figure 5**, respectively. Systems 3, 4A and 4B showed the highest fluorescence uniformity, with almost the entire field of view being at least 60% of the maximum fitted fluorescence intensity. In contrast, systems 1 and 2 showed lower uniformity, with less than half of the field of view reaching the 60% threshold. **Figure 4** shows the fitted fluorescence uniformity profiles and iso-maps for the systems, generated using the RBF interpolation method.

**Figure 4.**
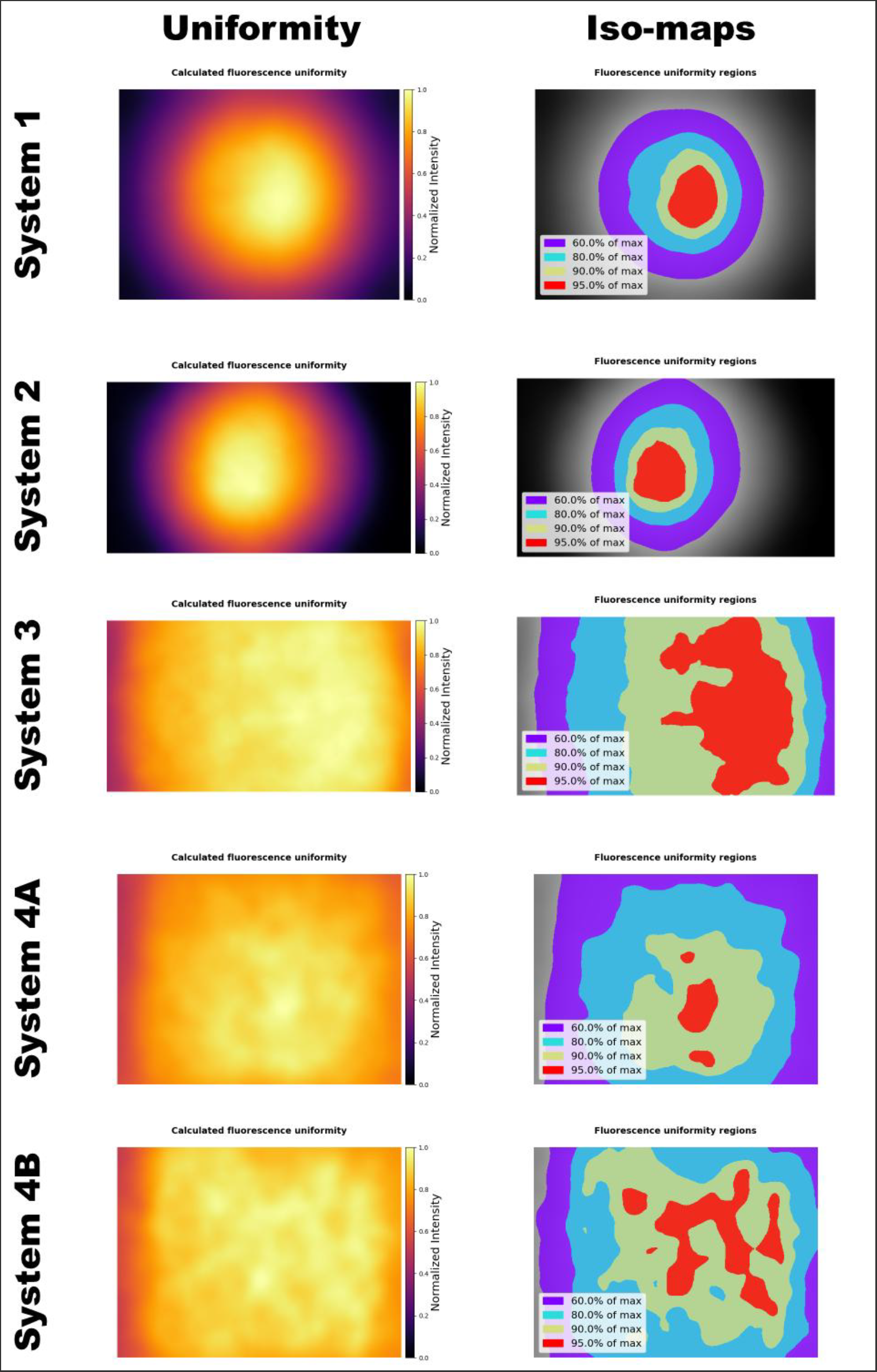
Fluorescence uniformity across the field of view of four commercial fluorescence imaging systems. Systems 3, 4A and 4B are the most uniform, with almost the entire field of view being at least 60% of the maximum fluorescence intensity. Systems 1 and 2 have much smaller regions where fluorescence intensity is at least 60% of maximum.

**Figure 5.**
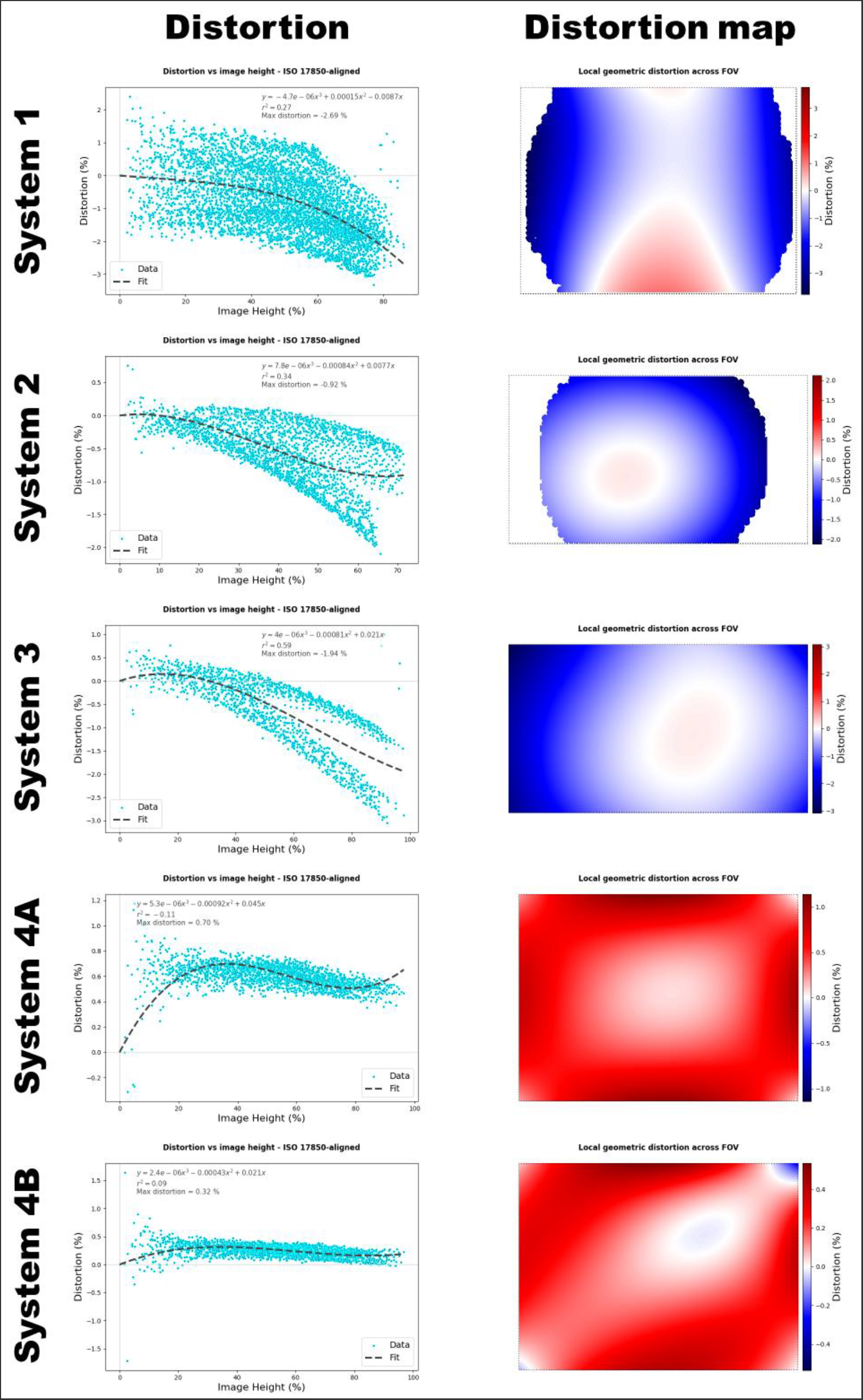
Local geometric distortion across the field of view of four commercial fluorescence imaging systems. System 1 has the most distortion, with a maximum of close to -3%. Systems 4A and 4B have the least distortion of less than 1%.

In terms of distortion, Systems 1, 2, and 3 showed some degree of overall negative distortion, while Systems 4A and 4B had minimal positive distortion. System 1 displayed the greatest distortion, approaching -3%, and had an asymmetric hourglass-like distortion pattern, where localized regions near the top and bottom center of the field of view showed slightly positive distortion. Systems 4A and 4B had the least distortion, both below 1%. The distortion graphs are shown in **Figure 5.**

### 3.4 Flat-field correction: RUD target

A total of 16 images were used to generate a fluorescence uniformity profile for the QUEL Imaging Box using the RBF interpolation method – the target was imaged at four locations spanning the field of view, in four orientations. The inverse of the resulting profile was multiplied by the input images to produce flat-field-corrected images. An example of one image pre and post this correction is shown in **Supplementary Figure S6**. These corrected images were then fed back into the custom Python code for assessing uniformity, and the resulting surface was flat within 95% of the max value. This is shown in **Supplementary Figure S7**. A second set of four images were acquired and these were corrected with the uniformity profile generated from the 16-image dataset. The resulting images were input to the uniformity assessment code and the results are shown in **Supplementary Figure S8**. The resulting uniformity profiles from these images are slightly less flat, however the entire field of view is again within 95% of the max value.

### 3.5 Flat-field correction: fluorescent cylinders

**Figure 6a,b** show the original images of the 100 nM and 0 nM fluorescent cylinders in the center and on the edge of the field of view. The corresponding images corrected with the uniformity profile from **Section 3.4** are shown in **Figure 6c,d**. In **Figure 6e,f**, the average intensities of the fluorescent cylinders are plotted in bar graphs. The measured ROI averages and standard deviations (fluorescence counts, arbitrary units) for the uncorrected 100 nM and 0 nM cylinders in the center of the field of view were 8976 ± 326 and 744 ± 37, respectively. The uncorrected values on the edge of the field of view were 6570 ± 277 and 636 ± 28. After flat-field correction, the ROI averages for the 100 nM and 0 nM cylinders were 9407 ± 354 and 764 ± 38 in the center of the field of view, and 9468 ± 558 and 952 ± 74 on the edge of the field of view. After correction, the intensity of the 100 nM well on the edge of the field of view matches its intensity in the center of the field of view. However, after correction, the intensity of the control (0 nM) on the edge of the field of view is greater than its intensity in the center. **Figure 7** shows a similar set of figures for the 3 nM and 0 nM pairing. Due to the low fluorescence intensity, the influence of the flat-field correction on the non-fluorescent background is more noticeable – the left and right edges of the corrected images have higher intensity than the center. In **Figure 7e,f**, the average intensities of both the 3 nM and 0 nM cylinders on the edge of the field of view overshoot their intensities in the center when the images are corrected.

**Figure 6.**
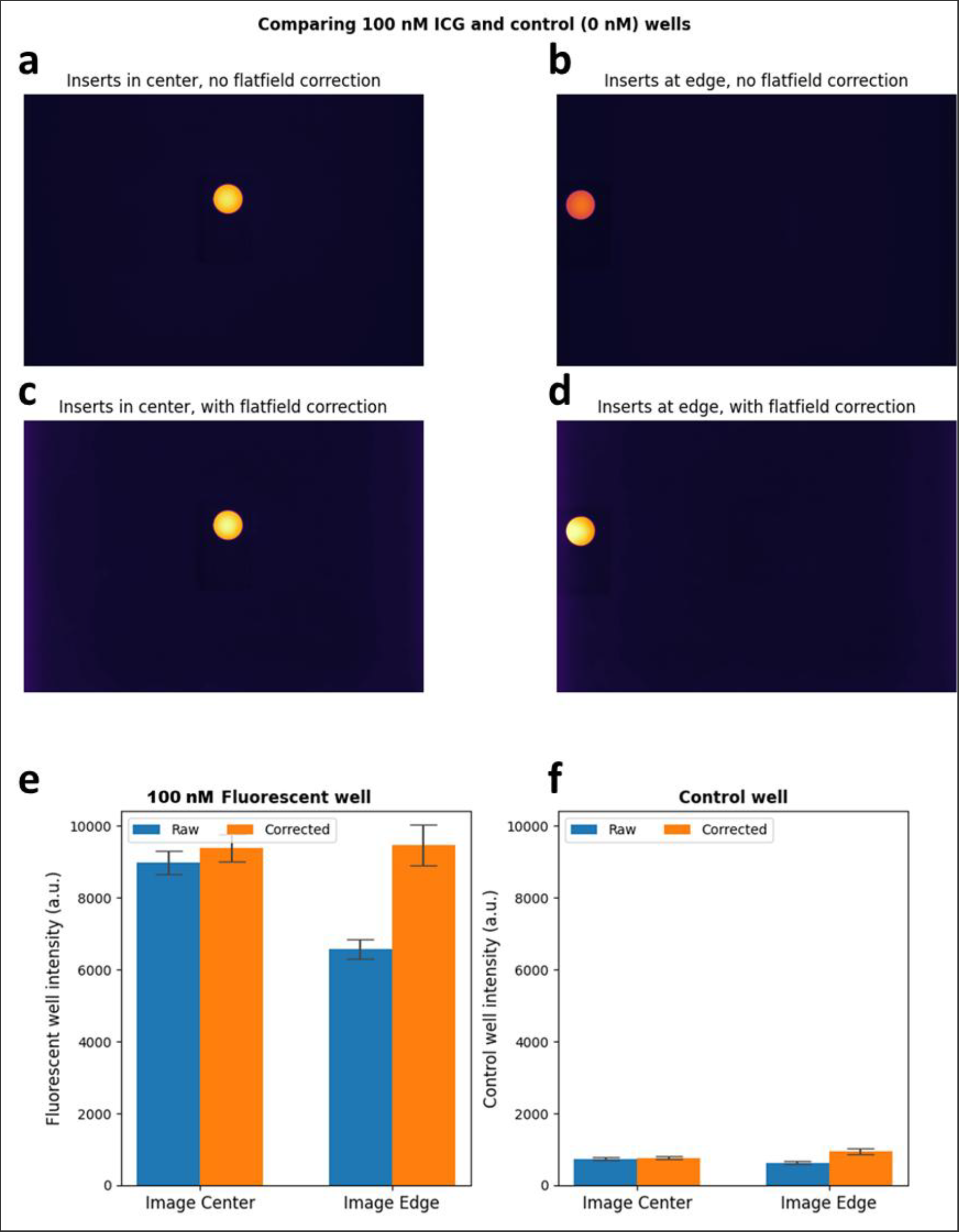
Comparison of 100 nM and 0 nM fluorescent cylinders with and without flat-field correction: shown are uncorrected images of the fluorescent cylinders in the center (a) and on the edge (b) of the field of view; followed by flat-field corrected images of the cylinders in the center (c) and on the edge (d) of the field of view; and finally, average intensities of an ROI centered on the 100 nM (e) and 0 nM (f) cylinder. When the image is flat-field corrected, the intensity of the 100 nM cylinder on the edge of the image matches its intensity in the center. However, the intensity of the control (0 nM) cylinder on the edge of the field of view overshoots the intensity in the center. Error bars in (e) and (f) depict ± one standard deviation.

**Figure 7.**
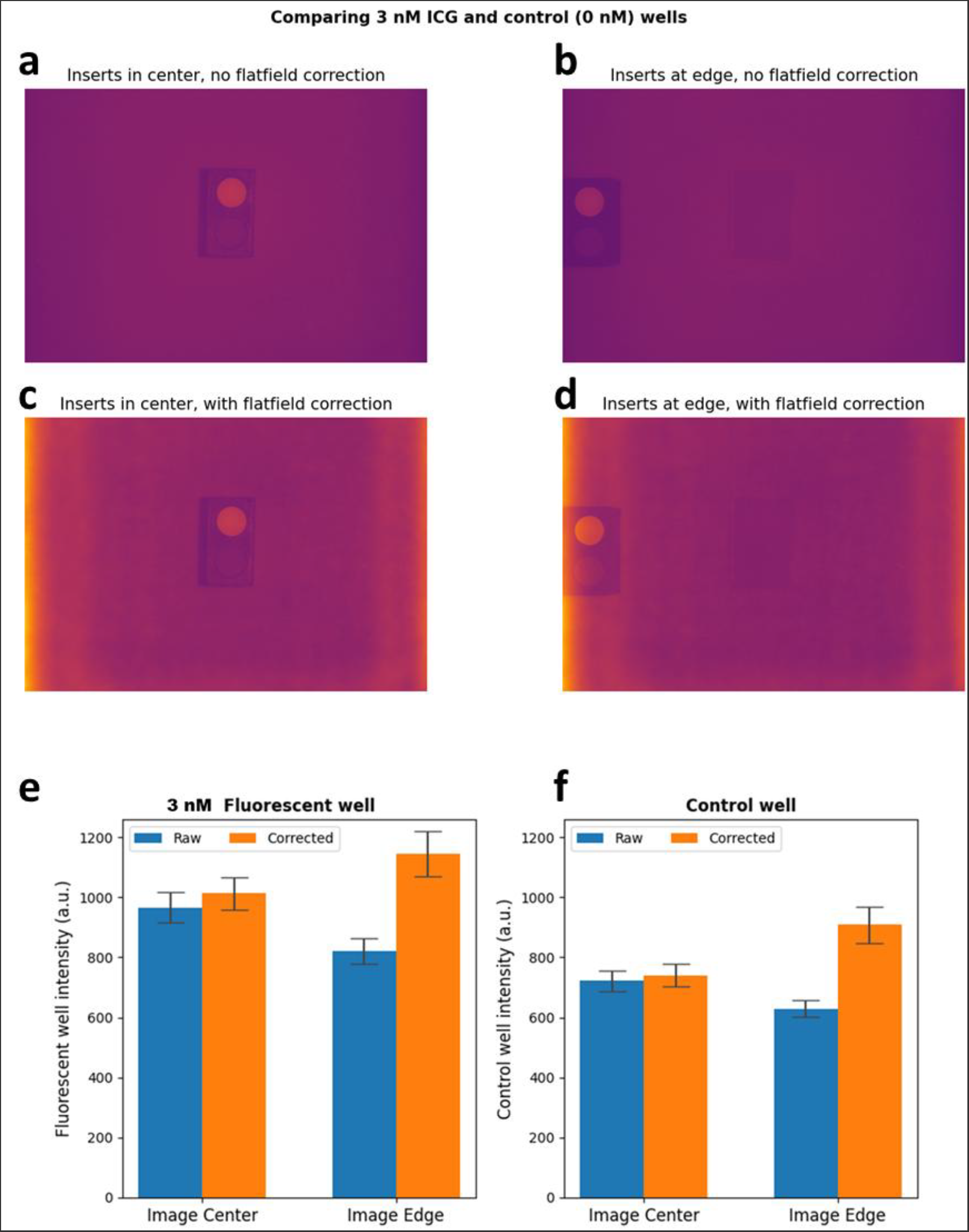
Comparison of 3 nM and 0 nM fluorescent cylinders with and without flat-field correction: shown are uncorrected images of the fluorescent cylinders in the center (a) and on the edge (b) of the field of view; followed by flat-field corrected images of the cylinders in the center (c) and on the edge (d) of the field of view; and finally, average intensities of an ROI centered on the 3 nM (e) and 0 nM (f) cylinder. When the image is corrected, the intensities of both the 3 nM and control (0 nM) cylinders on the edge of the field of view overshoot the corresponding intensities in the center. Error bars in (e) and (f) depict ± one standard deviation.

### 3.3 Flat-field correction: concentration sensitivity target

The acquired fluorescence images of the RCS target are shown in **Supplementary Figure S9**. Visually, flat-field correction improves the apparent consistency of fluorescence brightness across the images. Quantitative analysis of the images, however, shows that flat-field correction generally does not improve the measure of linearity, *m*, and in some cases worsens it substantially (*m* should ideally be equal to 1). Particularly, the corrected *Edge 0°* and *Edge 180°* images provide worse linearity metrics. **Figure 8** shows these results in log-log plots of normalized baselined fluorescence intensity versus fluorophore concentration. Additionally, **Table 1** contains the values of *m* for all five images, before and after flat-field correction. These findings highlight important caveats—namely, potential amplification of background signals and signal-dependent nonlinearity when corrected levels diverge from calibration conditions—and are further addressed in the Discussion section.

**Table 1.** Measure of linearity for RCS target images with and without flat-field correction.

|  | *m* |  |  |  |  |
| --- | --- | --- | --- | --- | --- |
|  | *Center* | *Edge, 0°* | *Edge, 90°* | *Edge, 180°* | *Edge, 270°* |
| **Original** | 1.010 | 1.070 | 0.997 | 0.939 | 0.897 |
| **Corrected** | 1.011 | 0.894 | 1.008 | 1.144* | 0.934 |

\* – For this image, the corrected control well had higher intensity than the corrected 1 nM well, resulting in negative intensity for the 1 nM well after baselining. Hence it was excluded from the fit to calculate $m$ .

**Figure 8.**
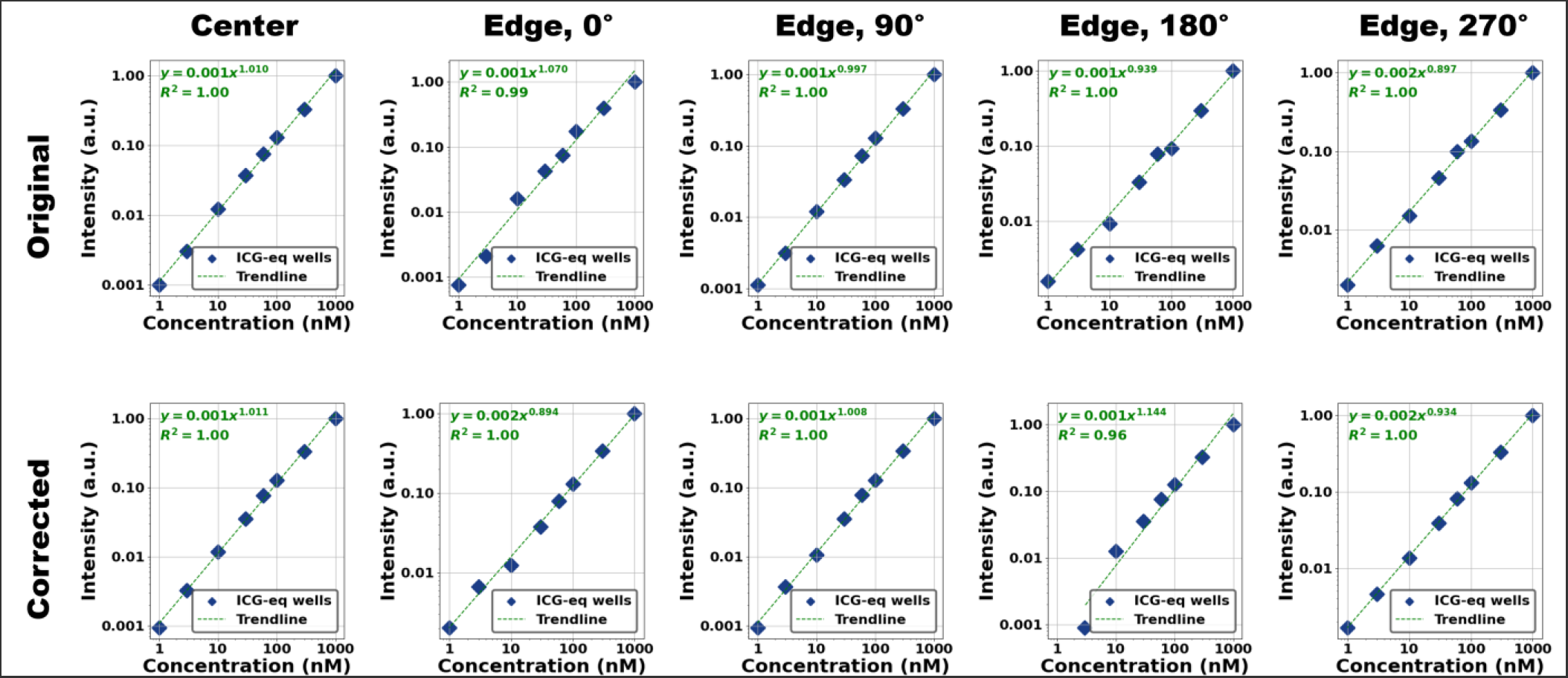
Results from analyzing images of the RCS target pre-(top row) and post-(bottom row) flat-field correction. Each graph is a log-log plot of normalized baselined fluorescence intensity versus fluorophore concentration. Flat-field correction does not generally improve the linearity fit, and in the case of the 180° rotation, substantially worsens it.

## 4 Discussion

Understanding the spatial uniformity of fluorescence detection across the field of view of a fluorescence imaging system is crucial, especially when clinical decisions depend on fluorescence intensity. Without this characterization, there is a risk of misinterpreting fluorescence images – such as deciding to perform unnecessary additional surgery based on perceived inadequate tissue perfusion during ICG angiography, or failing to identify fluorescently labeled cancerous tissue due to its position in a low-sensitivity region of the imaging field. It is also important to understand the spatial distortion of the imaging system, in particular for cases where the operator may not have visual access to the imaging subject, such as in endoscopic imaging.

While the uniformity of fluorescence capture is largely dependent on the illumination source, it is also affected by the spatial light collection efficiency of the optical assembly that detects the signal (i.e., relative illumination). Therefore, to truly assess fluorescence uniformity, simply mapping out the illumination profile, for example, with an optical power meter, is not sufficient. A homogeneously-fluorescent phantom, such as a bath of Intralipid into which fluorophore is diluted, can help capture the fluorescence uniformity profile of an imaging system.^14,33^ However, it is not shelf-stable, and can suffer from lack of repeatability. Moreover, the risk of contaminating the imaging area with fluorophore is high, and it cannot measure distortion. To address these limitations, we designed the RUD target, a solid phantom that contains a grid of equally fluorescent photostable wells that enables the characterization of both fluorescence uniformity and local geometric distortion simultaneously.

In this work, the RUD phantom was first evaluated on a custom fluorescence imaging system (QUEL Imaging Box) to demonstrate its ability to characterize fluorescence uniformity and geometric distortion. The analysis results generated with the QUEL-QAL library showed that this system has a rectangular-shaped fluorescence capture profile, with almost the entirety of the field of view being at least 60% of the maximum fluorescence intensity. Using the same image dataset, the geometric distortion across the imaging field was also assessed, revealing mild barrel distortion of less than 1%. Although our distortion analysis is consistent with ISO 17850:2015 standards, strict compliance with this standard requires specific conditions, including precise target alignment and dot size relative to the imaging field. Additionally, ISO 17850:2015 was originally developed for standard reflectance-based imaging (primarily white-light systems), and the use of image stitching for fluorescence systems with larger fields of view further deviates from this standard. It is worth noting that precise lateral displacement of the RUD between acquisitions allows subsampling of the spatial resolution (1 mm Ø wells with 2 mm spacing), effectively increasing the apparent density of the fluorescent dot-matrix when multiple images are combined.

Five commercial fluorescence imaging systems were characterized for fluorescence imaging uniformity and geometric distortion using the RUD target, demonstrating the phantom’s versatility across diverse device configurations. The results highlighted the differences in uniformity between these systems, with Systems 1 and 2, both surgical microscopes, exhibiting much smaller regions of fluorescence uniformity. However, these results are context-dependent: surgical microscopes typically utilize zoomed-in fields of view, focusing primarily on the image center; in contrast, the images used here were acquired with the field of view fully zoomed out. Improved uniformity for zoomed-in fields of view for surgical microscopes has been previously reported.^32^ System 3 is an exoscope, where fluorescence uniformity is likely more critical.^25^ System 4 is a small-animal imaging box, which would also require higher uniformity across the entire field-of-view. System 5 was unable to be evaluated due to the excitation wavelength of 805 nm, which exceeds the RUD target’s excitation range of 400-790 nm; future work may involve developing a version of the RUD target compatible with such systems. Distortion was generally small for these commercial systems, with System 1 showing the greatest distortion of ∼3%.

Using knowledge of an imaging system’s fluorescence uniformity profile to “flat-field” images is of interest in many applications, to improve qualitative appearance of the images and allow for quantitative analyses.^26–28^ In this paper, we demonstrated that the RUD target can be used for this application, by using it to capture the fluorescence uniformity profile of the QUEL Imaging Box, and then flat-field correcting a second set of images of the same target. As shown in **Supplementary Figures S6 – S8**, the corrected initial and second set of images had uniform fluorescence across the entire field of view (well within 95% of the maximum). Nevertheless, subsequent experiments caution the use of fluorescence flat-field correction for quantitative analyses.

In one experiment, a fluorescent 3D-printed cylinder containing 100 nM of ICG-equivalent fluorophore was imaged alongside a control (0 nM) cylinder in the center, then on the edge of the field of view of the QUEL Imaging Box. This 100 nM fluorescent cylinder had similar fluorescence signal intensity to the RUD target wells when imaged, and therefore when flat-field correction was performed, the corrected intensity on the edge of the image matched the corrected intensity in the center. In contrast, the corrected intensity of the control (which had no fluorophore) was higher on the edge of the field of view than in the center, highlighting inappropriate amplification of non-fluorescent signals. A similar effect was observed with a lower concentration (3 nM) cylinder, again suggesting incorrect intensity scaling resulting from the flat-field correction.

Further testing with the concentration sensitivity target, to span fluorescence intensities across multiple ICG-equivalent concentrations, showed that flat-field correction improved the linearity of imaged intensity vs. fluorophore concentration in only one of five imaging scenarios. In two scenarios, the flat-field correction substantially worsened linearity. As observed in the previous experiment, when the fluorescent intensity being corrected is similar to that of the RUD target, flat-field correction produces good results.

To understand why flat-field correction may negatively impact quantitative analyses, several factors must be considered. At low fluorophore concentrations, fluorescence intensity typically increases proportionally with concentration;^29^ however, practical imaging systems inherently have a detection threshold and a non-zero noise floor,^7^ below which proportionality no longer holds. Ideally, regions with no fluorophore should yield zero fluorescence intensity; however, autofluorescence and system background signals contribute to a baseline intensity that is always present, making it impossible to achieve zero intensity without artificial manipulation such as thresholding or baselining. Consequently, flat-field scaling factors derived from fluorescent signals can inadvertently amplify non-fluorescent background and autofluorescence signals disproportionately,^14^ as observed in our experiments with both fluorescent cylinders and the concentration sensitivity target. These observations are similarly noted in prior work emphasizing the limitations of flat-fielding for quantitative interpretation in intraoperative imaging.^14,33^ Additionally, at higher fluorophore concentrations, effects such as self-quenching can reduce fluorescence efficiency^30^, further disrupting the linear relationship between concentration and detected intensity. Thus, fluorescence flat-field correction is only reliable under conditions where fluorophore concentration and excitation intensity linearly correlate with the detected fluorescence signal, and background signals such as autofluorescence remain negligible.

Considering these limitations, extreme caution should be taken when using flat-field correction in quantitative analyses of *in vivo* fluorescence images, particularly because fluorophore concentration, autofluorescence, and background signal variations across the imaging field are typically unknown. For example, in ICG angiography where fluorescence is used as a measure of tissue perfusion,^31^ a piece of tissue on the edge of the field of view might appear dim because it has very little fluorophore (hence, perfusion), or because of the non-uniformity of the imaging system (needing flat-field correction). Hence, we recommend fluorescence flat-field correction only be performed for qualitative improvement of *in vivo* images. For quantitative assessment, it is better to identify the region of the imaging system’s field of view that is within an acceptable level of fluorescence uniformity, and only perform assessments within this region, which is consistent with prior work emphasizing the clinical impact of spatial non-uniformity and recommending center-region analysis for reliability.^32^ The RUD target and associated analysis code introduced in this manuscript can effectively facilitate this identification.

## 5 Conclusion

We developed a solid, shelf-stable fluorescent phantom (RUD) and accompanying open-source image analysis software (QUEL-QAL), for characterizing fluorescence uniformity and geometric distortion in fluorescence imaging systems. The utility of the RUD phantom and analysis code was successfully demonstrated on a custom-built imaging device and four commercial fluorescence imaging systems. The feasibility of fluorescence flat-field correction using RUD-generated uniformity profiles was also explored. Although flat-field correction qualitatively improved fluorescence image uniformity, it simultaneously can negatively impact quantitative accuracy. Consequently, we advise caution when applying flat-field correction in quantitative imaging applications and, instead of relying on this approach, recommend using uniformity assessment to identify regions of the field of view that meet acceptable uniformity criteria for quantitative measurements.

## Code and Data Availability

The data used in this study are available from the corresponding author upon reasonable request. The code used for the fluorescence uniformity and geometric distortion analysis is part of an open-source library, QUEL-QAL, developed to facilitate the analysis of fluorescence reference targets,^19^ available at: https://github.com/QUEL-Imaging/quel-qal (PyPI: quel-qal).

## Acknowledgements

This project was funded in part with federal funds from National Institute of Biomedical Imaging and Bioengineering, National Institutes of Health, Department of Health and Human Services, under grant number R43/44EB029804.

Acquisition of the Modus X exoscope (Synaptive Medical, Toronto, Ontario, Canada) at Dartmouth Hitchcock Medical Center was funded by National Institutes of Health: S10 OD032308-01.

