## Supplementary Figures for "A Phantom for Fluorescence Uniformity and Distortion Assessment of Near-Infrared Fluorescence Guided Surgery Systems"

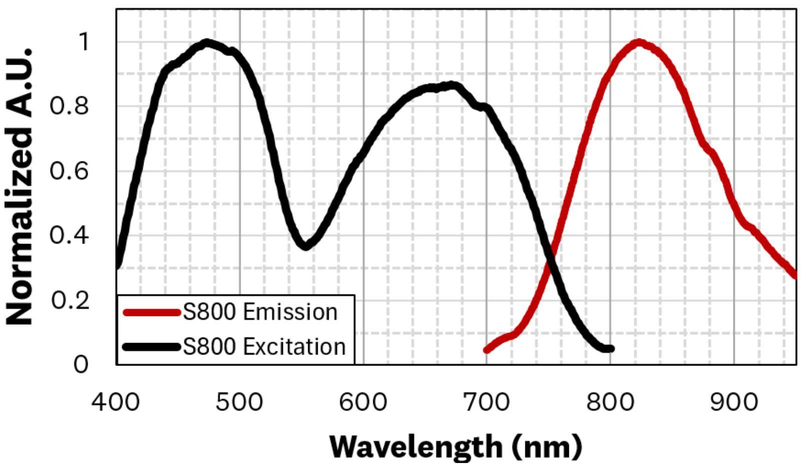


**Figure S1.** Excitation and emission spectra of the S800-01 luminescent compound

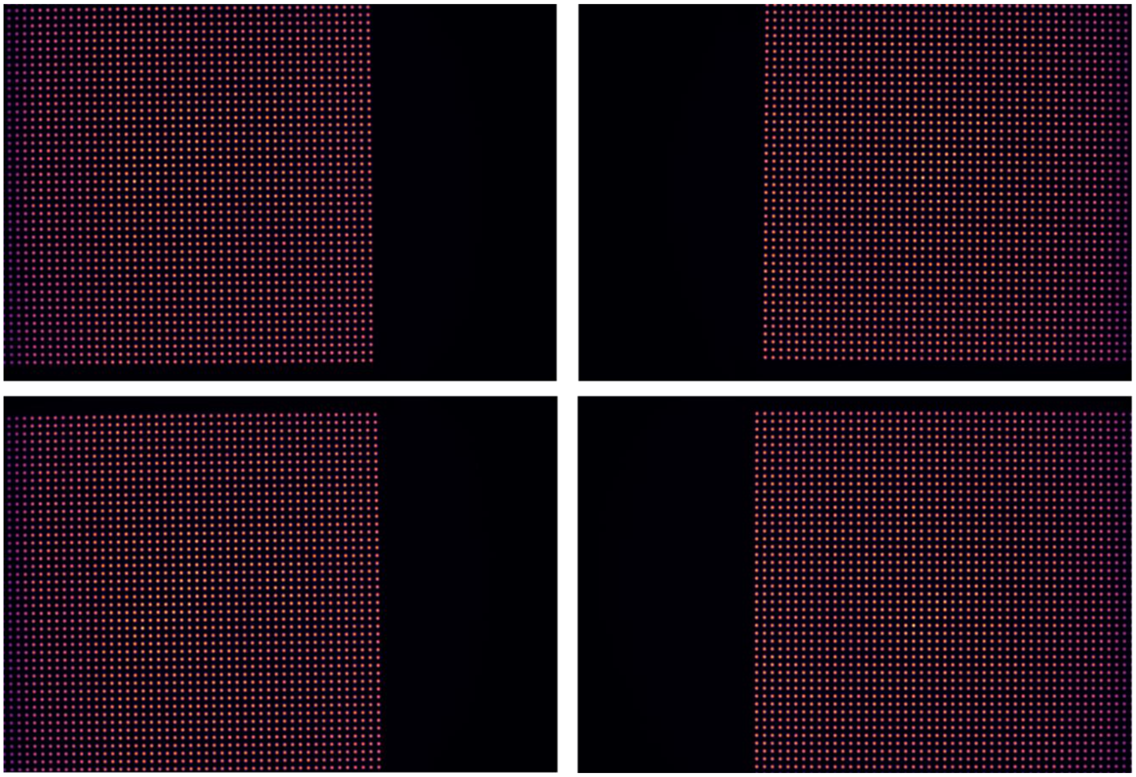

**Figure S2.** Four fluorescence images of the RUD target, taken so that data from the fluorescent wells span the entire field of view of the imaging system being evaluated.

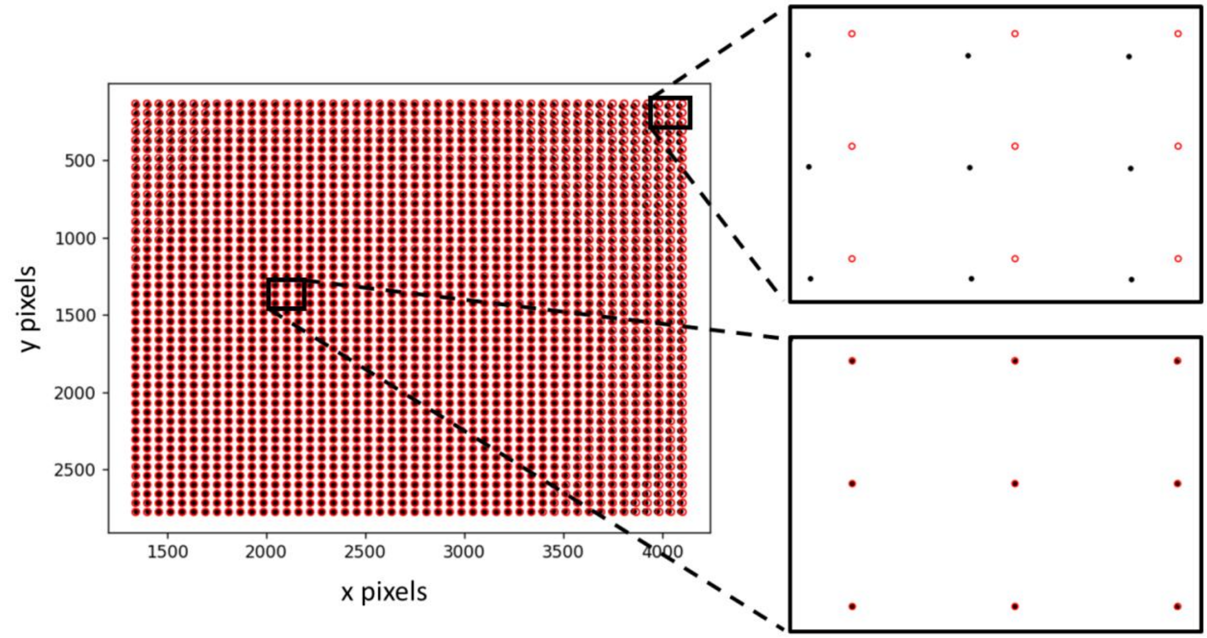


**Figure S3.** Visualization comparing the imaged well centroids (black circles) to the expected positions (red circles) derived from a regular reference grid. Towards the center of the image, the two overlap, but towards the periphery of the image, they diverge from each other.

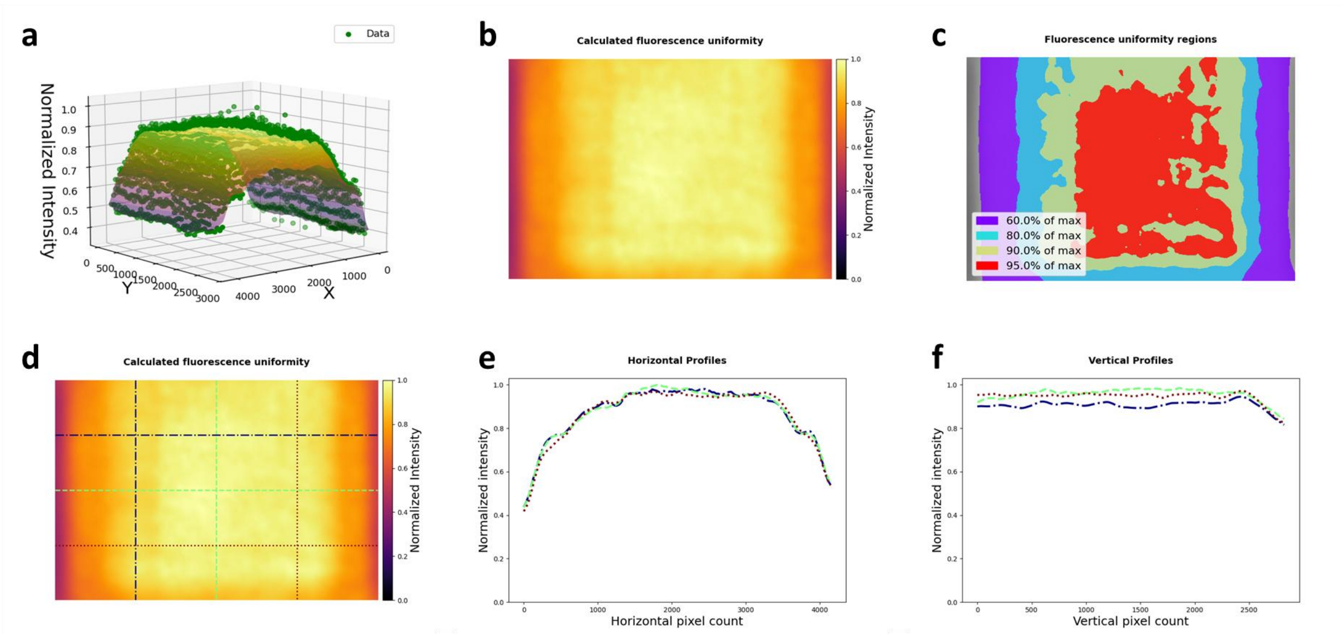


**Figure S4.** RUD analysis results using RBF interpolation method: (a) 3D plot showing extracted data and fitted surface; (b) fitted fluorescence uniformity map normalized to its maximum; (c) iso-maps, showing regions of the field of view that are at least 60%, 80%, 90%, and 95% of the maximum intensity; (d – f) line profiles across the fluorescence uniformity fit. Note the transient dips in intensity on the outer edges of the uniformity profile that are not present in the results of the b-spline fit.

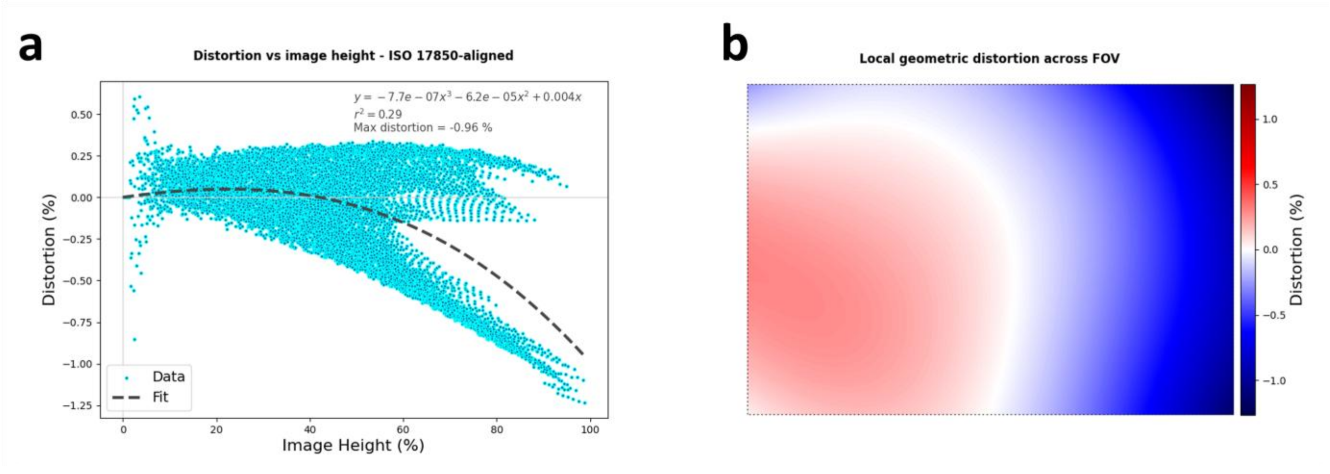


**Figure S5.** Distortion analysis results after placing 3° wedge underneath RUD target: (a) local geometric distortion as a function of image height, showing two groups of data points, one with slightly positive, and the other with more negative distortion; (b) spatial map of distortion across the field of view, showing clear keystone distortion where the left side of the image has positive distortion and the right side of the image has negative distortion. FOV = field of view.

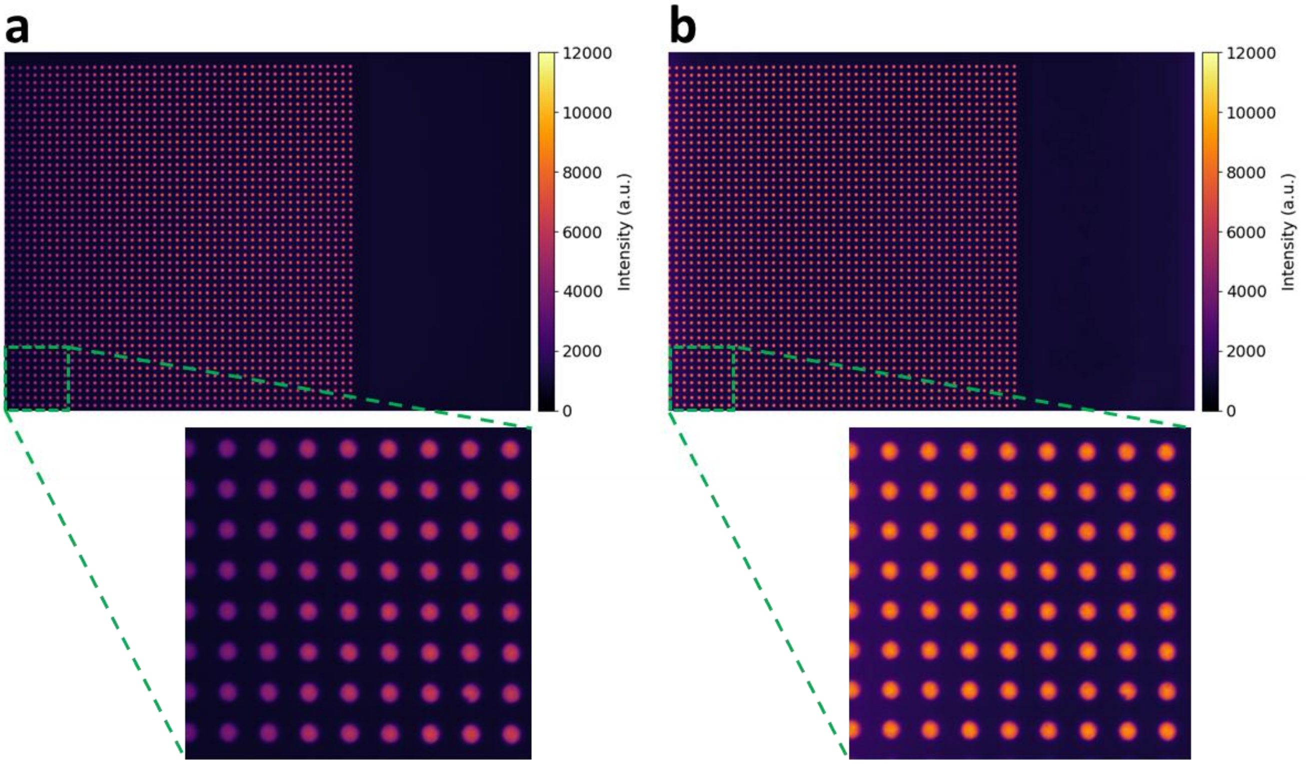


**Figure S6.** A fluorescence image of the RUD target pre- (a), and post- (b) flatfield correction. The fluorescent wells are more uniform in intensity after the correction.

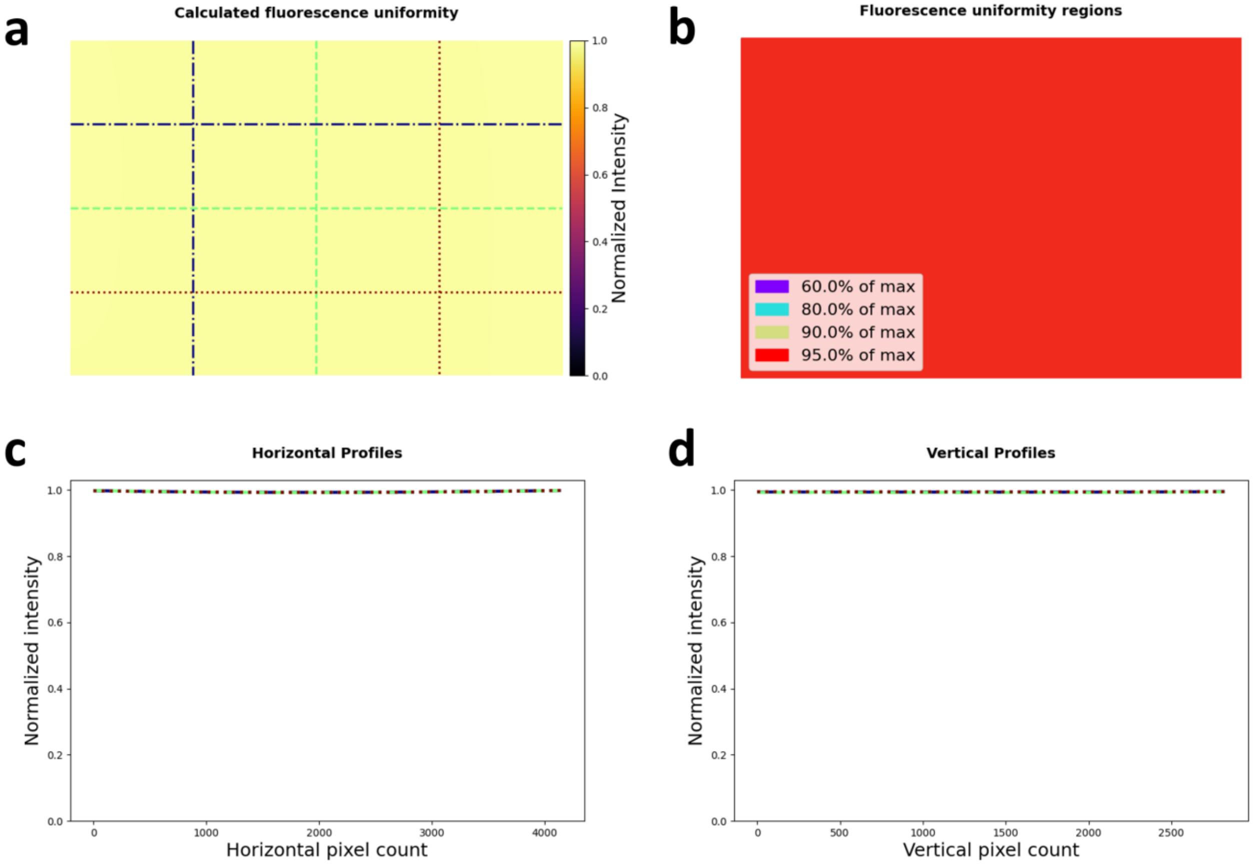


**Figure S7.** Results from uniformity analysis of flat-field corrected RUD images: line profiles are visually very flat (a, c, d); iso-maps, showing that the entire field of view is at least 95% of the max intensity (b).

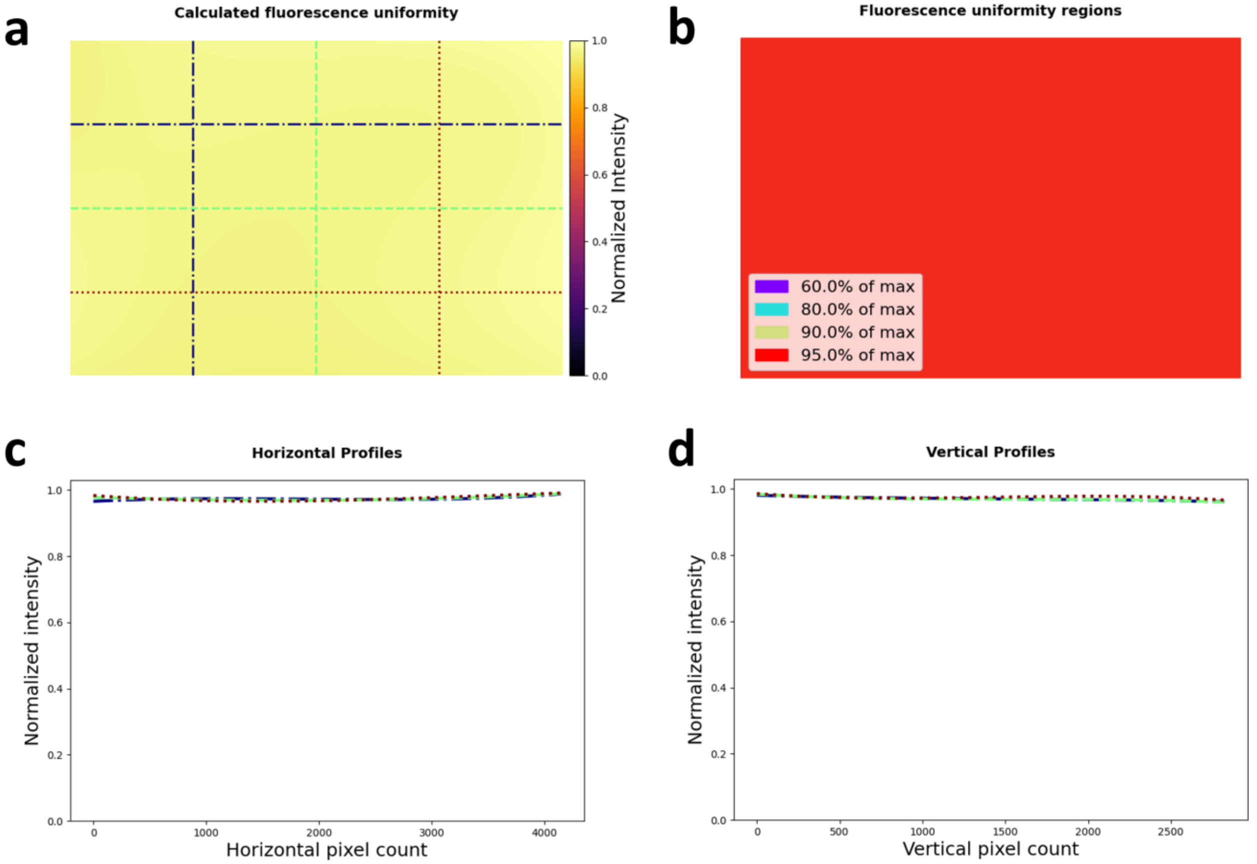


**Figure S8.** Results from uniformity analysis of new set of flat-field corrected RUD images that were not used in generating the applied uniformity profile: line profiles are a little wavier (a, c, d); however, iso-maps show that the entire field of view is still at least 95% of the max intensity (b).

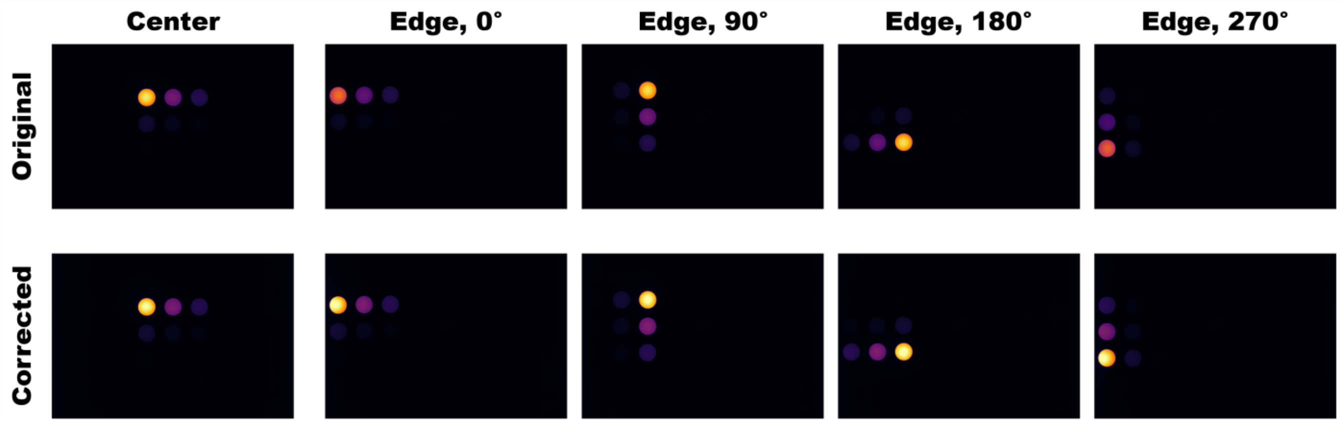


**Figure S9.** Fluorescence images of the RCS target pre- (top row) and post- (bottom row) flatfield correction. When corrected, the images appear more consistent regardless of orientation of the RCS target or location within the field of view.
